# Genomic Characterization of Lung Cancer in Never-Smokers Using Deep Learning

**DOI:** 10.1101/2025.08.14.670178

**Authors:** Monjoy Saha, Thi-Van-Trinh Tran, Praphulla MS Bhawsar, Tongwu Zhang, Wei Zhao, Phuc H. Hoang, Karun Mutreja, Scott M. Lawrence, Nathaniel Rothman, Qing Lan, Robert Homer, Marina K. Baine, Lynette M. Sholl, Philippe Joubert, Charles Leduc, William D. Travis, Stephen J. Chanock, Jianxin Shi, Soo-Ryum Yang, Jonas S. Almeida, Maria Teresa Landi

## Abstract

Despite promising results in using deep learning to infer genetic features from histological whole-slide images (WSIs), no prior studies have specifically applied these methods to lung adenocarcinomas from subjects who have never smoked tobacco (NS-LUAD) – a molecularly and histologically distinct subset of lung cancer. Existing models have focused on LUAD from predominantly smoker populations, with limited molecular scope and variable performance. Here, we propose a customized deep convolutional neural network based on ResNet50 architecture, optimized for multilabel classification for NS-LUAD, enabling simultaneous prediction of 16 molecular alterations from a single H&E-stained WSI. Key architectural modifications included a simplified two-layer residual block without bottleneck layers, selective shortcut connections, and a sigmoid-based classification head for independent prediction of each alteration, designed to reduce computational complexity while maintaining predictive accuracy. The model was trained and evaluated on 495 WSIs from the Sherlock-*Lung* study (70% training with 10% internal test set for 10-fold cross-validation, and 30% held-out validation set for final evaluation). For the held-out validation data, our model achieved high areas under the receiver operating characteristic curve [AUROC] values =0.84-0.93 for detecting 11 features: *EGFR, KRAS, TP53, RBM10* mutations, *MDM2* amplification, kataegis, *CDKN2A* deletion, *ALK* fusion, whole-genome doubling, and *EGFR* hotspot mutations (p.L858R and p.E746_A750del). Performance was low to moderate for tumor mutational burden (AUROC=0.67), APOBEC mutational signature (AUROC=0.57), and *KRAS* hotspot mutations (p.G12C: AUROC=0.74, p.G12V: AUROC=0.55, p.G12D: AUROC=0.43). Compared to results from established architectures such as Inception-v3 on the same WSIs, our model demonstrated significantly improved performance for most features. With further optimization, our model could support triaging for molecular testing and inform precision treatment strategies for NS-LUAD patients.

## Introduction

Lung cancer in never-smokers (LCINS), defined as individuals who have never smoked or smoked less than 100 cigarettes in their lifetime, accounts for 10–25% of lung cancer cases [1]. Considered separately, LCINS represents the fifth leading cause of cancer-related mortality globally [2]. LCINS is predominantly classified as adenocarcinoma (here, never-smoker lung adenocarcinoma, NS-LUAD) and presents distinct clinical, histological, and molecular characteristics [1, 2]. LCINS is characterized by a high prevalence of targetable driver mutations, such as in *EGFR* (identified in 34–55% of cases vs. 7–18% in smoking related cases), and *ALK* (rearrangements,4–9% vs. 2–3%) [2-4].

Molecular testing—especially through next-generation sequencing (NGS)—has become standard in lung cancer care, primarily to identify targetable driver mutations for which effective targeted therapies are available. NGS also enables simultaneous profiling of co-occurring genomic alterations—both actionable and non-actionable—offering a more comprehensive view of tumor biology. This expanded molecular landscape is critical for developing combination strategies aimed at improving clinical outcomes and overcoming therapeutic resistance [5, 6]. However, despite its clinical value, the broad implementation of NGS remains limited by practical constraints, including diagnostic turnaround time, tissue availability, and cost especially in resource-limited settings.

In this context, machine learning (ML) and deep learning (DL) models offer a scalable, tissue-sparing, and cost-efficient alternative for predicting molecular alterations directly from hematoxylin and eosin (H&E)-stained whole-slide images (WSIs), a resource that is readily available in routine pathology workflows [7]. Proof-of-concept studies—mostly trained on highly curated public datasets such as The Cancer Genome Atlas or single-institution datasets—have employed convolutional neural networks (CNNs), including different variants of ResNet [8] and Inception [9] networks, to predict molecular alterations from WSIs [10-17]. However, most existing models have focused on a limited subset of molecular features (e.g., *EGFR, KRAS*, and *ALK* mutations), while other clinically relevant alterations, including hotspot gene variants, copy number alterations, and global genomic characteristics remain largely unexplored, limiting the potential of DL models as comprehensive alternatives for broad-panel NGS. Previously reported performances varied considerably, with area under the curve (AUROC) values ranging from 0.45 to 0.92, and cross-institutional generalization remains a significant challenge, reflecting the limitations of single-institution designs and the limited diversity in patient demographics, tissue quality, and imaging protocols [11-15, 18]. Moreover, previous models were trained with a poor representation of never-smokers, raising concerns about generalizability to LCINS.

In this study, we proposed a customized deep convolutional neural network—referred to as the customized multilabel CNN—built upon the ResNet50 [8] architecture. The model was trained and evaluated using NS-LUAD cases from the Sherlock-*Lung* study, a large, diverse, international, multi-institutional cohort [19]. In our held-out validation set, the model can simultaneously predict accurately 11 genomic alterations from H&E-stained WSIs, including nonsynonymous mutations, hotspot driver mutations including different genotypes within the same gene, gene fusions, copy number alterations, mutational signatures, and global molecular features. This approach provides a robust, scalable framework for image-based and genotype-specific molecular characterization of NS-LUAD that could be deployed in clinical settings.

## Materials and methods

### Sherlock-Lung whole-slide image dataset

The ongoing Sherlock-*Lung* study, initiated in 2019, recruited confirmed treatment-naïve, NS-LUAD patients from multiple institutions in Asia, Africa, North America, South America, Europe, and Australia [19]. The study obtained comprehensive clinical data from medical records, tumor specimens, and H&E-stained WSIs. Tumor samples underwent whole-genome sequencing (WGS) as previously described [4, 19, 20]. WSIs were digitized using image scanners from various manufacturers, including Hamamatsu and Leica.

For this study, we obtained clinical data from medical records provided by participating centers, including age at tumor diagnosis, sex, tumor stage (American Joint Committee on Cancer 8th edition), and histologic grade (2020 International Association for the Study of Lung Cancer system) [21]. We assigned genetic ancestry using WGS data based on genetic similarity to 1000 Genomes Project (1KGP) “super population” European, African, East Asian and Ad Mixed American reference samples via VerifyBamID2 [20].

Of 552 WSIs from 467 individual cases, 57 were excluded from analysis because of the absence of mutations in 15 of 16 profiled molecular features of interest (excluding tumor mutational burden [TMB]). After exclusions, the final dataset included 495 WSIs from 410 cases, including 424 (85.7%) formalin-fixed and 71 (14.3%) frozen samples. Diagnosis and morphological features were confirmed by at least two lung pathologists for each case.

### Image preprocessing pipeline

The WSI dataset was randomly divided into training (*n* = 346) and held-out validation (*n* = 149) sets, with each patient contributing WSIs exclusively to one set. Within the training set, 10% of the slides (n = 35) were reserved for stratified ten-fold cross-validation. The training set was used for model training, the internal test set (from stratified ten-fold cross-validation) was used for hyperparameter tuning, and the held-out validation set was reserved exclusively for final model evaluation. To evaluate differences in baseline clinical characteristics and molecular feature distributions between the two cohorts, we used Student’s t-test for continuous variables, and the χ^2^ test for categorical variables. For categorical variables with fewer than five WSIs in any category, we used Fisher’s exact test.

Non-overlapping 256 × 256-pixel tiles were extracted at 20× magnification using the OpenSlide Python library, yielding a total of 5,316,629 tiles—3,935,357 from the training set (including internal test tiles used in ten-fold cross-validation) and 1,381,272 from the held-out validation set reserved for final evaluation. We did not perform region-level annotations (e.g., tumor vs. adjacent normal tissue); however, most WSIs predominantly contained tumor regions. To minimize noise during model training, tiles containing less than 50% tissue were excluded from the training set. Labels for molecular alterations were assigned at the patient level, without spatial annotation. Image preprocessing involved rescaling pixel values to the [0,1] range, along with extensive data augmentation during training. Augmentation strategies included random rotations, width and height shifts, and horizontal and vertical flips, aiming to increase data diversity and mitigate overfitting.

Sixteen molecular features were assigned to each WSI (**Table 1**), including nonsynonymous and driver mutations, fusions, and copy number alterations in known lung cancer driver genes (i.e., *TP53, KRAS, EGFR, CDKN2A, MDM2, ALK, RBM10*); APOBEC mutational signature; kataegis; whole-genome doubling (WGD); TMB; and hotspot mutations in *EGFR* (p.L858R, p.E746_A750del) and *KRAS* (p.G12C, p.G12V, p.G12D). Besides the known targetable mutations, the selection of these features was guided by the availability of data, existing research gaps, and previous studies from our group in LCINS [4, 22]. All features were dichotomized: driver mutations as wild-type or mutant, and all other features as absent or present, except for TMB, which was dichotomized at the median value (median: 1.83; range: 0.28–22.33).

**Table 1.** Clinical and molecular characteristics of never-smokers lung adenocarcinomas.

|  | Total<br>(n=495 WSIs)<br>N (%) | Training + Internal testing<br>(cross-validation) set<br>(n=346 WSIs)<br>N (%) | Held-out<br>validation set<br>(n=149 WSIs)<br>N (%) | p-value |
| --- | --- | --- | --- | --- |
| **Age at Diagnosis (years): Mean (Median, Range)** | 65 (66, 26 - 90) | 65 (66, 26 - 90) | 65 (67, 39 - 90) | 0.98 |
| **Sex** |  |  |  |  |
| Male | 116 (23) | 81 (23) | 35 (23) | 1.00 |
| Female | 379 (77) | 265 (77) | 114 (77) |  |
| **WGS-based ancestry** |  |  |  |  |
| European | 332 (67) | 236 (68) | 96 (64) | 0.71 |
| African | 2 (0) | 1 (0) | 1 (1) |  |
| East Asian | 143 (29) | 98 (28) | 45 (30) |  |
| Admixed American | 18 (4) | 11 (3) | 7 (5) |  |
| **AJCC 8<sup>th</sup> edition stage** |  |  |  |  |
| I | 266 (54) | 185 (53) | 81 (54) | 0.88 |
| II | 74 (15) | 52 (15) | 22 (15) |  |
| III | 97 (20) | 67 (19) | 30 (20) |  |
| IV | 28 (6) | 22 (6) | 6 (4) |  |
| Unknown | 30 (6) | 20 (6) | 10 (7) |  |
| **IASLC classification** |  |  |  |  |
| Grade 1 | 94 (19) | 68 (20) | 26 (17) | 0.72 |
| Grade 2 | 149 (30) | 107 (31) | 42 (28) |  |
| Grade 3 | 55 (11) | 35 (10) | 20 (13) |  |
| Unclassified | 197 (40) | 136 (39) | 61 (41) |  |
| **Driver Mutations<sup>1</sup>** |  |  |  |  |
| EGFR | 218 (44) | 149 (43) | 69 (46) | 0.63 |
| EGFR p.L858R | 99 (20) | 72 (21) | 27 (18) | 0.53 |
| EGFR p.E746_A750del | 71 (14) | 45 (13) | 26 (17) | 0.27 |
| KRAS | 92 (19) | 62 (18) | 30 (20) | 0.69 |
| KRAS p.G12C | 24 (5) | 19 (5) | 5 (3) | 0.42 |
| KRAS p.G12D | 27 (5) | 18 (5) | 9 (6) | 0.89 |
| KRAS p.G12V | 31 (6) | 17 (5) | 14 (9) | 0.10 |
| TP53 | 143 (29) | 101 (29) | 42 (28) | 0.85 |
| RBM10 | 40 (8) | 28 (8) | 12 (8) | 1.00 |
| **Other molecular features<sup>1</sup>** |  |  |  |  |
| APOBEC | 181 (37) | 126 (36) | 55 (37) | 0.95 |
| CDKN2A deletion | 86 (17) | 61 (18) | 25 (17) | 0.78 |
| MDM2 amplification | 71 (14) | 56 (16) | 15 (10) | 0.06 |
| ALK fusion | 49 (10) | 31 (9) | 18 (12) | 0.27 |
| WGD | 131 (26) | 97 (28) | 34 (23) | 0.19 |
| Kataegis | 127 (26) | 93 (27) | 34 (23) | 0.36 |
| TMB: Mean (Median, Range) | 2.45 (1.83, 0.28 – 22.33) | 2.49 (1.81, 0.28-22.33) | 2.34 (1.88, 0.31-12.11) | 0.57 |

<sup>1</sup>The percentage of slides with a specific molecular alteration. AJCC: American Joint Committee on Cancer, IASLC: International Association for the Study of Lung Cancer, TMB: Tumor mutational burden, WGD: Whole-genome doubling, WSI: Whole slide image

### Molecular and Genomic Characterization with Customized Deep Learning Architectures: Multilabel and Binary Convolutional Neural Networks

We developed a customized ResNet50 [8] architecture designed to reduce computational complexity while maintaining the core principles of residual learning. The network begins with a 7×7 convolutional layer with 64 filters, followed by batch normalization, ReLU activation, and a max pooling layer. The body of the architecture is composed of custom residual blocks that preserve the foundational ideas of ResNet, including skip connections and identity mappings. However, unlike the original ResNet50 which employs three-layer bottleneck blocks (1×1, 3×3, 1×1), our custom design simplifies each residual block to two convolutional layers, both using a consistent 3×3 kernel size. This design choice maintains representational power while reducing parameter count and computational overhead. Each residual block includes two sequential convolutional layers with batch normalization and ReLU activation, followed by an additive skip connection. When the input and output dimensions differ due to stride variations, a 1×1 convolution is applied to the shortcut path to ensure dimensional consistency. This conditional shortcut mechanism ensures that residual learning is preserved across all stages of the network. Following the residual blocks, the feature maps undergo global average pooling, which reduces spatial dimensions and preserves salient features. This is followed by a fully connected layer with 100 ReLU-activated units, a dropout layer (rate = 0.5) to prevent overfitting, and a final output layer with sigmoid activation.

In the multilabel classification setting, the output layer comprises 16 sigmoid-activated units, each representing the probability of a distinct molecular alteration. The model was trained using the binary cross-entropy loss function applied independently to each output node. This architecture provides a balance between model capacity and efficiency, making it well-suited for biomedical datasets where data volume is typically limited. A schematic overview of the proposed workflow architecture is provided in **Fig. 1**. The code is publicly available (see Code Availability).

**Fig. 1.**
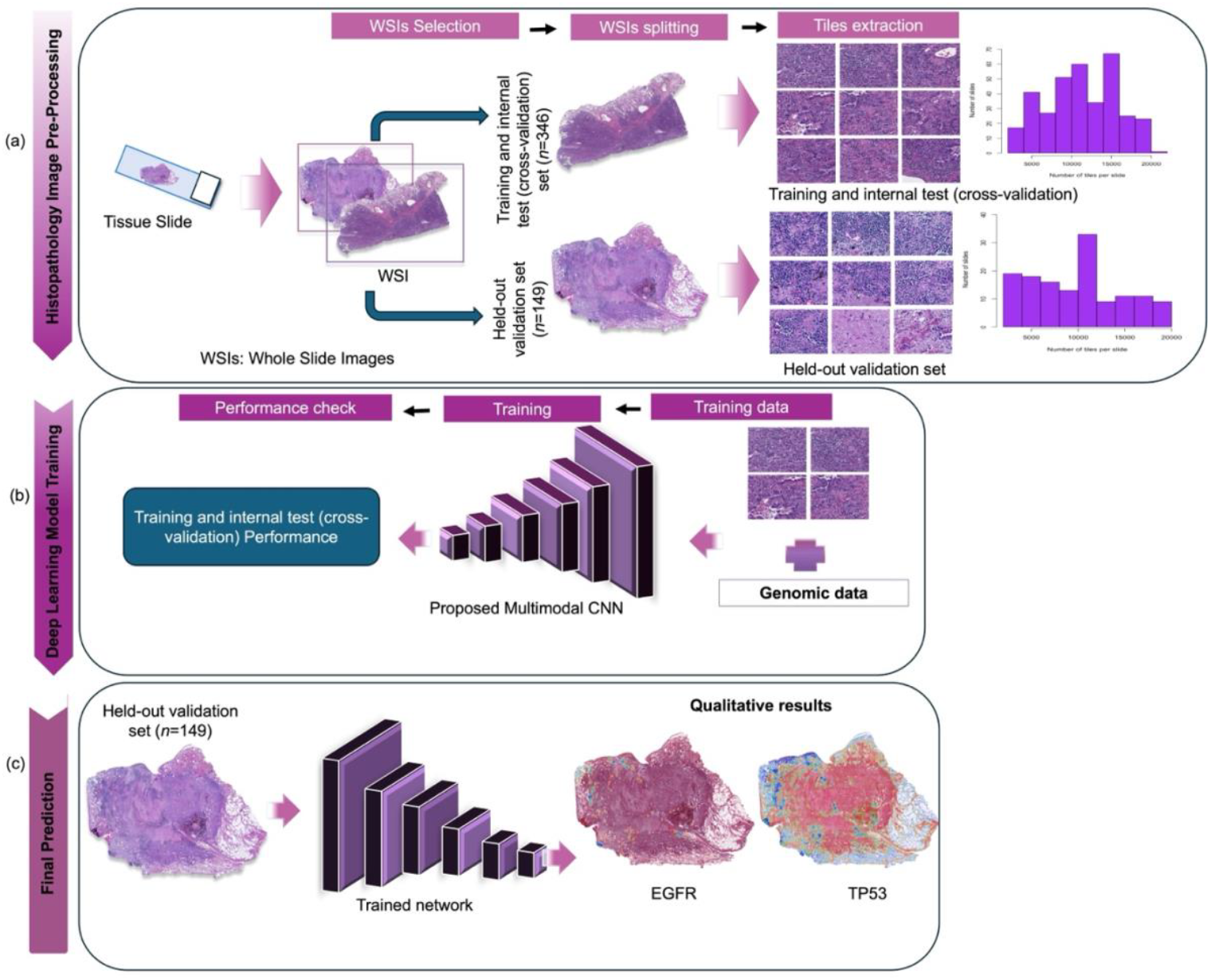
Overview of the workflow for image-based molecular prediction in NS-LUAD. The diagram illustrates the end-to-end pipeline for histopathology (a) image preprocessing, (b) deep learning model training, and (c) molecular prediction. (a) Tissue slides were digitized into WSIs, which were split into training and internal test (cross-validation) set (n = 346), and a held-out validation set (n = 149). Within the training set (n = 346), 10% of the slides (n = 35) were reserved as an internal test set for stratified ten-fold cross-validation. Image tiles were extracted from each set, with accompanying bar charts showing tile distribution. (b) A customized convolutional neural network was trained using paired image and genomic data (i.e., nonsynonymous mutations, gene fusions, and copy number alterations in key driver genes (TP53, KRAS, EGFR, CDKN2A, MDM2, ALK, RBM10), APOBEC mutational signature, and other molecular features [kataegis, whole-genome doubling, tumor mutational burden], and hotspot mutations in EGFR [p.L858R, p.E746_A750del] and KRAS [p.G12C, p.G12V, p.G12D]). (c) Model performance was assessed on the held-out validation set. The training, internal test (cross-validation), and held-out validation datasets were constructed to be mutually exclusive. Heatmaps were generated using Gradient-weighted Class Activation Mapping, with representative heatmaps shown for predictions of EGFR and TP53 mutations.

For comparison, we developed a binary classification CNN, using the same architecture but reconfiguring it with a single sigmoid-activated output node to predict the wild-type or mutant status, or the presence or absence of a specific molecular or genomic characteristic. Of note, we employed the same training, internal test (cross-validation), and held-out validation sets for both the multilabel and binary approaches. Although this binary approach eliminates inter-label competition and allows for focused training on individual labels, it required training 16 separate models to cover all target alterations, resulting in a significantly higher computational cost compared to the single multilabel model.

### Training, Internal testing (cross-validation) and Validation Procedure

The training, internal testing (cross-validation), and validation procedures for the multilabel and binary classification models were designed to be consistent and comparable, differing only in output structure, batch sampling and evaluation metrics. Both models used binary cross-entropy loss, appropriate for both multilabel (independent binary outputs) and binary classification tasks. The training, internal testing (cross-validation), and validation datasets were constructed to be mutually exclusive. Both the training and internal testing (cross-validation) phases utilized image data in combination with gene and other genomic variables, while the validation phase relied solely on image data. Training was conducted using the Adam optimizer with a learning rate of 0.0001 and a batch size of 64. Momentum was not applied explicitly, consistent with the Adam optimizer’s design. Contrary to an earlier grid search plan, hyperparameters such as learning rate, batch size, and optimizer choice were standardized across models to isolate the effect of output structure rather than optimization configuration. Additionally, early stopping was implemented with a patience of 15 epochs, and model checkpointing was used to preserve the best model based on validation binary accuracy. Stratified ten-fold cross-validation was applied to evaluate model performance reliably across both binary and multilabel classification settings, using AUC as the performance metric. Given the varying prevalence of molecular features in our dataset— from common (e.g., *EGFR* mutations, 44%; APOBEC mutational signature presence, 37%; *TP53* mutations, 29%) to rare (e.g., *KRAS* p.G12C hotspot mutation, 5%; *ALK* fusion, 10%)— stratification was essential to preserve label distribution across folds. For multilabel classification, batch sampling was adapted to balance classes more evenly, improving learning on rare labels. During inference, optimal classification thresholds were computed per label using Youden’s index, maximizing the sum of sensitivity and specificity. This per-label threshold tuning helped mitigate bias from imbalanced label distributions by tailoring decision boundaries to each class. In contrast, the binary classification model used a single global threshold, also derived through Youden’s index, simplifying the decision rule for the single output. In the binary setting, stratification maintained balanced representation of positive and negative classes for each molecular feature, while for multilabel classification, a multilabel-stratified approach ensured that the co-occurrence and frequency of multiple labels per sample were proportionally represented in each fold. This approach mitigated sampling bias, ensured representative learning subsets, and supported robust model training and fair comparison across all folds.

### Evaluation metrices

To assess the performance of our customized multilabel CNN model, we assessed its ability to simultaneously predict genomic alterations per sample using a held-out validation dataset. We computed receiver operating characteristic (ROC) curves, and corresponding AUROC values. We estimated 95% confidence intervals for AUROC values using 1000 bootstrap resampling iterations [23, 24]. We also calculated sensitivity, specificity, and Youden’s index [25]. Youden’s index, defined as sensitivity + specificity − 1, summarizes diagnostic accuracy, with values ranging from 0 (no discrimination) to 1 (perfect performance). The optimal classification threshold was determined by maximizing Youden’s index.

We compared our customized multilabel CNN model with established deep learning architectures— Inception-V3 [15] and standard ResNet50 [13, 26] on the held-out validation set. Differences in predictive performance, as assessed by ROC curves, were evaluated using the DeLong method [27]. Multiple comparisons were adjusted using the Benjamini–Hochberg correction method [28]. All tests were two-sided, with p-value < 0.05 considered statistically significant. Statistical analyses were conducted using the scipy.stats module in Python.

### Generation of activation scores and heatmaps

We applied the Gradient-weighted Class Activation Mapping (Grad-CAM) technique [29, 30] to produce activation scores over the WSIs, highlighting class-discriminative spatial regions. Activation scores were derived by multiplying each pixel of the input images with the corresponding weights from the final convolutional layer of the customized Multilabel CNN, ranging from 0 (no activation) to 1 (strong activation). These activation scores represent the model’s confidence that a given tissue region contributes to the prediction of a specific molecular feature. The resulting scores were visualized using the different colormaps (e.g., Jet) from Python libraries to depict the spatial localization of predictive signals.

The heatmaps were generated without pixel-or region-level annotations, using only slide-level molecular labels. This weakly supervised learning paradigm allows the model to infer spatial coordinates associated with specific molecular features without explicit distinction between tumor and normal tissue. The resulting maps highlight predictive regions irrespective of cellular identity. **Supplementary Fig. 1** shows a screenshot of our interactive web-based tool that displays activation heatmaps for a representative case of lung adenocarcinoma with a *TP53* mutation in an individual who has never smoked. The heatmaps were visualized using the *OpenSeadragon* viewer [31] integrated with the *ImageBox3* tiling mechanism [32], enabling interactive exploration of high-resolution WSIs.

## Results

### Cohort’s Clinical and Molecular Characteristics

Subjects’ demographic and clinical characteristics as well as prevalence of tumor molecular features are described in **Table 1**. Briefly, the median age at diagnosis was 65 years (range: 26–90), and 77 % (*n*=379) of cases were female. Based on WGS-derived genetic ancestry, this cohort includes 332 (67%) patients of European ancestry from the United States, Canada, and Europe; 143 (29%) of East Asian ancestry from Hong Kong, Taiwan, the United States, and Canada; 18 (4%) of admixed American ancestry from Europe and Canada, and 2 (0.4%) of African ancestry from the United States and Canada. The prevalence of genomic alterations ranged from ∼5% for the *KRAS* p.G12D hotspot mutation to ∼44% for *EGFR* mutations consistent with prior observations. We observed no significant differences in clinical and molecular characteristics between the training/ Internal testing (cross-validation) and held-out validation sets.

### Evaluation of the customized multilabel CNN on the held-out validation set

We evaluated the performance of the customized multilabel model on the Sherlock-*Lung* held-out validation set, generating ROC curves for both driver mutations (**Fig. 2a**) and other molecular features (**Fig. 2b**). Corresponding AUROC scores with 95% confidence intervals are summarized in **Table 2**. The multilabel CNN model demonstrated high predictive accuracy (AUROC ≥ 0.8) for *EGFR* (AUROC = 0.93), *KRAS* (AUROC = 0.92), *TP53* (AUROC = 0.91), *RBM10* (AUROC = 0.90), *EGFR* p.L858R (AUROC = 0.92) and *EGFR* p.476_A750del (AUROC = 0.86) mutations. Performance was lower across *KRAS* hotspot mutations, with AUROC values of 0.74 for p.G12C, 0.55 for p.G12V, and 0.43 for p.G12D, likely because of their small sample size. For other molecular features, the model achieved strong performance for *MDM2* amplification (AUROC = 0.93), Kataegis (AUROC = 0.91), *CDKN2A* deletion (AUROC = 0.89), *ALK* fusion (AUROC = 0.86), and WGD (AUROC = 0.84), while showing lower performance for TMB (AUROC = 0.67) and APOBEC mutational signature (AUROC = 0.57).

**Table 2.** Comparing the performance of customized multilabel CNN with established deep learning architectures, Inception-V3 [15] and ResNet50 [13, 26], using AUROC and 95% confidence intervals [CI] on the held-out validation dataset (n=149). P-values are from DeLong’s [27] test between the customized multilabel CNN and other models.

|  | *Molecular alterations* | *Customized*<br>*Multilabel CNN*<br>*AUROC, 95% CI* | *Inception-V3*<br>*AUROC, 95% CI* | *p-value* | *ResNet50*<br>*AUROC, 95% CI* | *p-value* |
| --- | --- | --- | --- | --- | --- | --- |
| Driver Mutation | EGFR | 0.93<br>[0.90, 0.96] | 0.84<br>[0.79, 0.88] | **<0.01** | 0.91<br>[0.88, 0.94] | 0.38 |
|  | KRAS | 0.92<br>[0.89, 0.95] | 0.82<br>[0.77, 0.89] | **<0.01** | 0.89<br>[0.85, 0.93] | 0.24 |
|  | TP53 | 0.91<br>[0.87, 0.94] | 0.86<br>[0.82, 0.90] | **0.02** | 0.90<br>[0.86, 0.94] | 0.80 |
|  | RBM10 | 0.90<br>[0.86, 0.93] | 0.72<br>[0.66, 0.78] | **<0.01** | 0.84<br>[0.79, 0.88] | **0.03** |
|  | EGFR p.L858R | 0.92<br>[0.89, 0.95] | 0.75<br>[0.69, 0.80] | **<0.01** | 0.87<br>[0.83, 0.91] | **0.04** |
|  | EGFR p.E476_A750del | 0.86<br>[0.82, 0.90] | 0.70<br>[0.64, 0.76] | **<0.01** | 0.76<br>[0.71, 0.81] | **<0.01** |
|  | KRAS p.G12C | 0.74<br>[0.68, 0.80] | 0.63<br>[0.57, 0.69] | **<0.01** | 0.69<br>[0.63, 0.75] | 0.24 |
|  | KRAS p.G12V | 0.55<br>[0.48, 0.61] | 0.5<br>[0.43, 0.57] | 0.18 | 0.51<br>[0.44, 0.58] | 0.43 |
|  | KRAS p.G12D | 0.43<br>[0.37, 0.49] | 0.5<br>[0.40, 0.58] | 0.12 | 0.45<br>[0.38, 0.51] | 0.77 |
| Molecular Features | CDKN2A deletion | 0.89<br>[0.85, 0.93] | 0.82<br>[0.77, 0.87] | **<0.01** | 0.86<br>[0.80, 0.92] | 0.43 |
|  | MDM2 amplification | 0.93<br>[0.90, 0.96] | 0.77<br>[0.71, 0.82] | **<0.01** | 0.87<br>[0.81, 0.93] | 0.09 |
|  | ALK fusion | 0.86<br>[0.82, 0.90] | 0.69<br>[0.63, 0.75] | **<0.01** | 0.80<br>[0.73, 0.87] | 0.17 |
|  | WGD status | 0.84<br>[0.79, 0.88] | 0.81<br>[0.74, 0.89] | 0.39 | 0.84<br>[0.78, 0.90] | 1.00 |
|  | Kataegis | 0.91<br>[0.88, 0.94] | 0.87<br>[0.81, 0.93] | 0.18 | 0.90<br>[0.85, 0.95] | 0.80 |
|  | APOBEC | 0.57<br>[0.51, 0.63] | 0.55<br>[0.47, 0.63] | 0.59 | 0.57<br>[0.49, 0.65] | 1.00 |
|  | TMB | 0.67<br>[0.60, 0.73] | 0.63<br>[0.55, 0.71] | 0.33 | 0.66<br>[0.58, 0.74] | 0.90 |

CNN: Convolutional neural network, TMB: Tumor mutational burden, WGD: Whole-genome doubling, A two-sided $p < 0.05$ is considered as statistically significant.

**Fig. 2.**
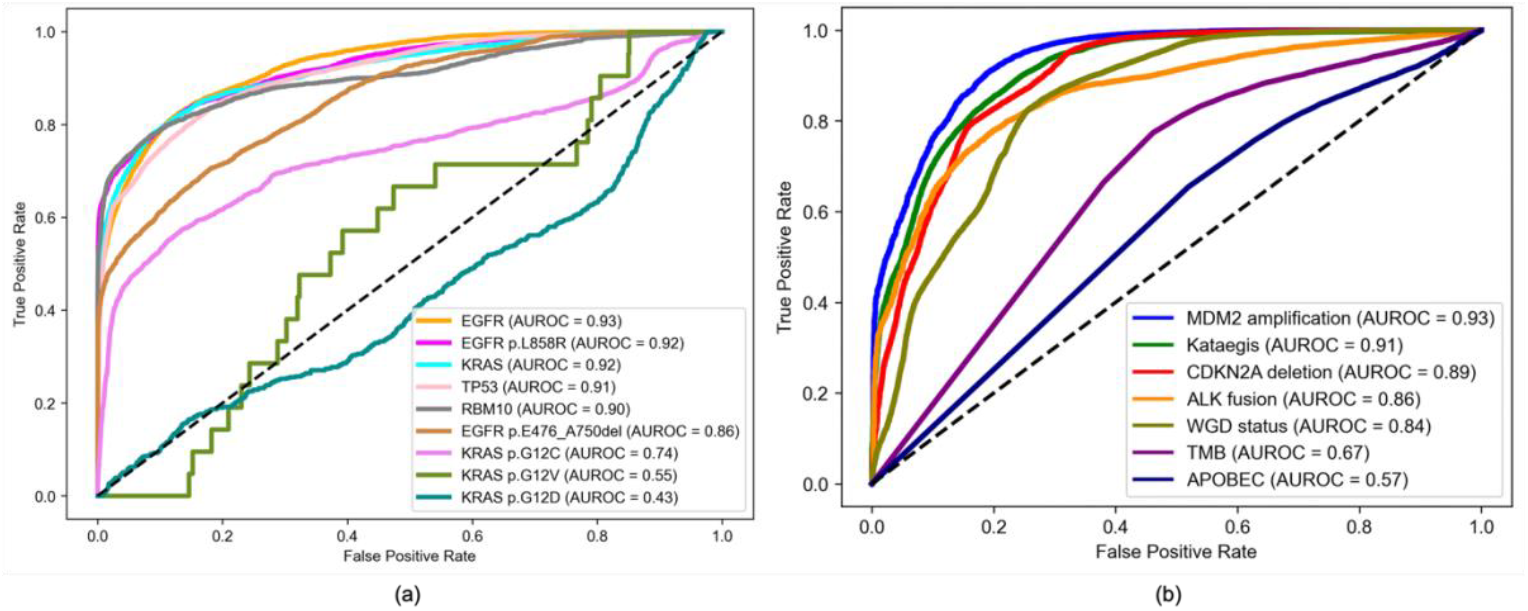
Receiver operating characteristic curves and corresponding area under the ROC curve (AUROC) values for the customized multilabel convolutional neural network evaluated on the held-out validation set (n = 149) of hematoxylin and eosin (H&E)-stained whole-slide images from the Sherlock-*Lung* cohort. (a) driver gene mutations and hotspot variants (*EGFR, KRAS, TP53*, and *RBM10, EGFR* p.L858R, *EGFR* p.E476_A750del, *KRAS* p.G12C, *KRAS* p.G12D, *KRAS* p.G12V) and for (b) other molecular features, including copy number alterations (*CDKN2A* deletion, *MDM2* amplification), gene fusions (*ALK*), and global genomic characteristics (whole-genome doubling, APOBEC mutational signature, tumor mutational burden). All predictions were generated using a single trained multilabel model. The order of alterations presented in the figure reflects descending AUROC values, from highest to lowest. TMB: Tumor mutational burden, WGD: Whole-genome doubling.

Sensitivity, specificity, and Youden’s Index for molecular alterations with AUROC ≥ 0.80 are shown in **Table 3**. Sensitivity was ≥0.80 for *EGFR, EGFR* p.L858R, and *TP53* mutations, *CDKN2A* deletion, *MDM2* amplification, WGD, and kataegis; and reached 0.78 and 0.71 for *KRAS* and *EGFR* p.E746_A750del mutations respectively. Specificity was ≥0.80 for *EGFR, EGFR* p.L858R, *EGFR* p.E476_A750del, *TP53, RBM10, KRAS* mutations, *CDKN2A* deletion, *MDM2* amplification, and *ALK* fusion and was lower for kataegis (0.76) and WGD status (0.74).

**Table 3.** Sensitivity, specificity, and Youden’s Index for molecular features with AUROC values of ≥0.8 using the customized multilabel CNN model on H&E-stained WSIs.

|  | Molecular alterations | Sensitivity | Specificity | Youden’s Index |
| --- | --- | --- | --- | --- |
| Driver Mutations | EGFR | 0.82 | 0.87 | 0.70 |
|  | EGFR p.L858R | 0.83 | 0.86 | 0.69 |
|  | EGFR p.E476_A750del | 0.71 | 0.82 | 0.52 |
|  | TP53 | 0.80 | 0.86 | 0.65 |
|  | RBM10 | 0.78 | 0.92 | 0.69 |
|  | KRAS | 0.78 | 0.91 | 0.69 |
| Molecular features | CDKN2A deletion | 0.80 | 0.83 | 0.63 |
|  | MDM2 amplification | 0.90 | 0.81 | 0.72 |
|  | ALK fusion | 0.78 | 0.80 | 0.58 |
|  | WGD status | 0.83 | 0.74 | 0.57 |
|  | Kataegis | 0.90 | 0.76 | 0.65 |

CNN: Convolutional neural network, WGD: Whole-genome doubling

### Comparison with binary and benchmark CNNs on the held-out validation set

Compared with independently trained binary CNNs, which are more computationally intensive, the customized multilabel model showed near-identical performance across all features (**Table 2, Figure 3)**, with no evidence of systematic bias, despite reduced computational complexity. Moreover, the customized multilabel CNN consistently outperformed Inception-V3 and standard ResNet50 in predicting molecular alterations, with statistically significant improvements for *EGFR* (Inception-V3: AUROC = 0.84), *KRAS* (Inception-V3: AUROC = 0.82,), *TP53* (Inception V3: AUROC = 0.86), *RBM10* (Inception V3: AUROC = 0.72, ResNet50: AUROC = 0.84), *EGFR* p.L858R (Inception-V3: AUROC = 0.75, ResNet50: AUROC = 0.87), *EGFR* p.E476_A750del (Inception-V3: AUROC = 0.70, ResNet50: AUROC = 0.76), *KRAS* p.G12C (Inception-V3: AUROC = 0.63), *CDKN2A* deletion (Inception-V3: AUROC = 0.82, ResNet50: AUROC = 0.86), *MDM2* amplification (Inception-V3: AUROC = 0.770, ALK fusion (Inception-V3: AUROC = 0.69,). For other molecular alterations, the customized multilabel CNN performed comparably or marginally better than Inception-V3 and ResNet50, although the differences were not statistically significant (**Table 2**).

**Fig. 3.**
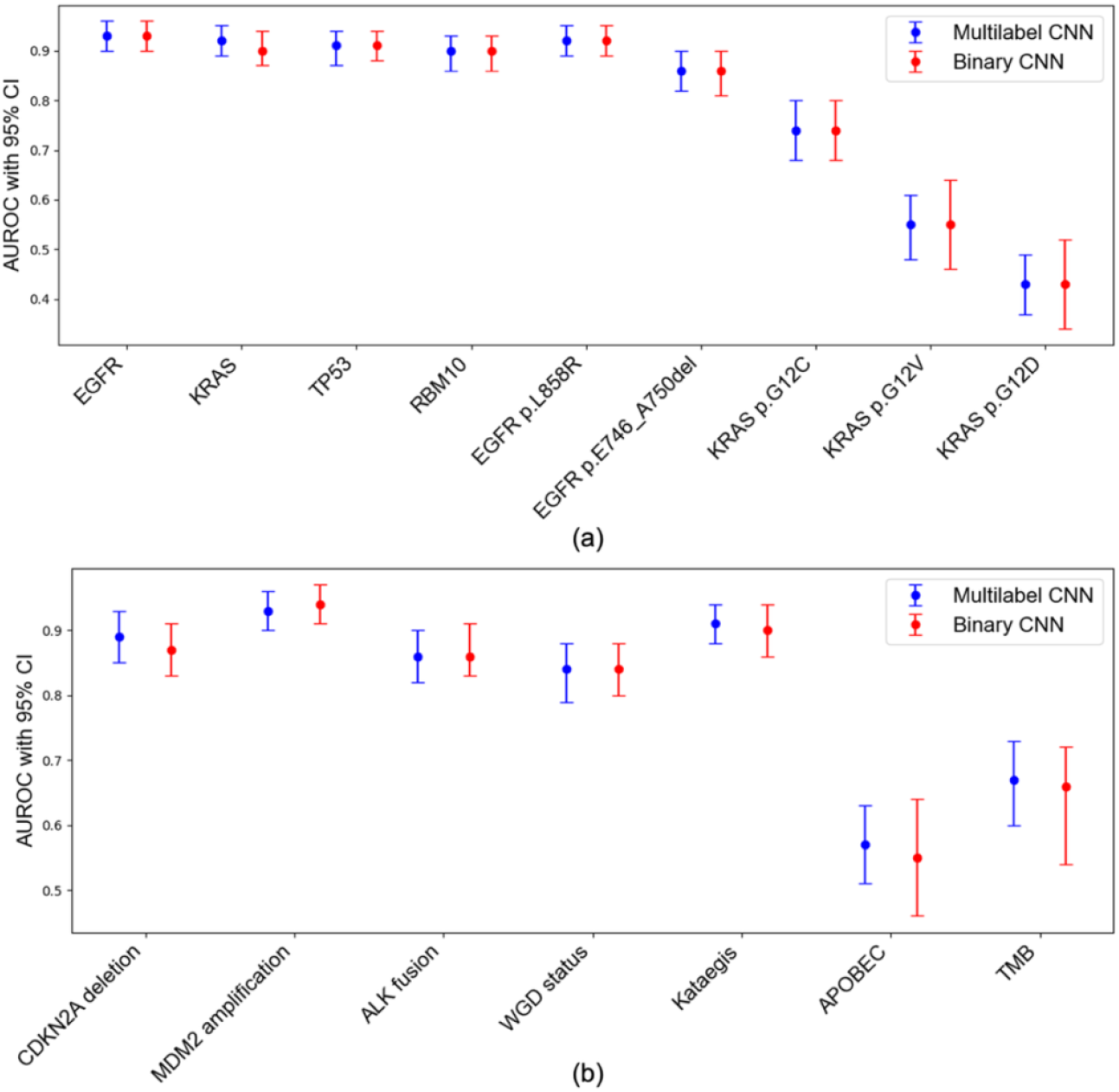
Comparison of predictive performance between multilabel and binary CNNs across molecular features. AUROC scores with 95% confidence intervals (CIs) for the multilabel (blue) and binary (red) CNN models. Each point represents the AUROC for a given feature, with error bars indicating 95% bootstrap confidence intervals. (a) Driver gene mutations and hotspot variants (EGFR, KRAS, TP53, and RBM10, EGFR p.L858R, EGFR p.E476_A750del, KRAS p.G12C, KRAS p.G12D, KRAS p.G12V). (b) other molecular features, including copy number alterations (*CDKN2A* deletion, *MDM2* amplification), gene fusions (*ALK*), and global genomic characteristics (whole-genome doubling, APOBEC mutational signature, tumor mutational burden). The multilabel model achieves comparable performance to the binary models across all features, despite reduced computational complexity.

### Reconstruction of predicted molecular features from customized multilabel CNN model

To interpret model predictions, we generated Grad-CAM heatmaps overlaid on WSIs of tumors with representative molecular profiles. **Figures 4a** and **4b** illustrate two examples, each demonstrating that our customized multilabel CNN accurately predicts multiple molecular alterations with spatially resolved activation patterns.

**Fig. 4.**
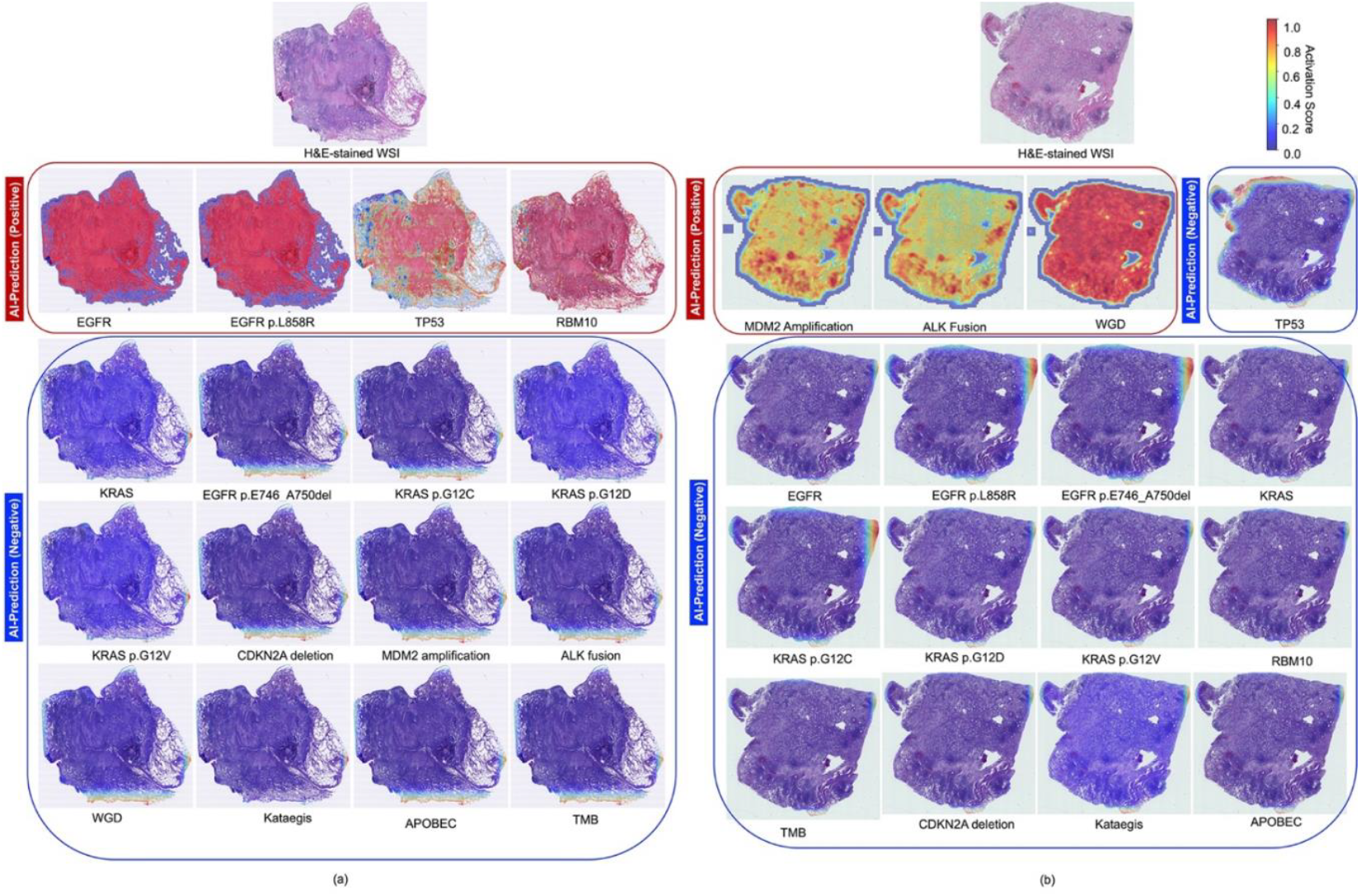
Representative activation heatmaps for predicted driver genes and molecular features in NS-LUAD, generated by our proposed multilabel convolutional neural network. Two held-out validation cases, randomly selected from the n=149 held-out validation set, are illustrated in panels (a) and (b). Each panel shows the original H&E-stained whole-slide image alongside the corresponding activation heatmaps for 16 molecular alterations. These heatmaps highlight either positive or negative predictions of key driver mutations and molecular features made by our multilabel CNN. In panel (a), the tumor harbors *EGFR, EGFR* p.L858R, *TP53*, and *RBM10* mutations. The model correctly predicts the presence of all four (indicated by red boxes), while the remaining 12 features—including *KRAS* variants, *CDKN2A* deletion, *MDM2* amplification, *ALK* fusion, whole-genome doubling, Kataegis, APOBEC, and tumor mutational burden —are predicted absent (blue boxes). In panel (b), the tumor exhibits *MDM2* amplification, *ALK* fusion, and WGD, all of which are accurately predicted, and the remaining molecular alterations are negative. Activation heatmaps are color-coded by confidence scores (red = high, blue = low), reflecting the model’s certainty for each prediction. Notably, these heatmaps were generated without pixel-level annotations, demonstrating the effectiveness of a weakly supervised training strategy. Red-highlighted regions indicate the areas the CNN deemed most informative for classifying each molecular feature. An interactive web-based platform was also developed to enable real-time visualization of these heatmaps as “digital stains” to assist pathologists in interpretation (see *Code Availability*). For details on the heatmap generation methodology, refer to *Methods*, section (vii).

**Fig. 4a** presents a WSI of a tumor harboring an *EGFR* mutation—specifically the hotspot p.L858R variant—alongside *TP53* and *RBM10* mutations, confirmed by WGS. The corresponding heatmaps display strong activation in tumor regions associated with these alterations predicted by the customized multilabel CNN, with minimal activation across the remaining molecular features. Similarly, **Fig. 4b** shows a tumor with confirmed *MDM2* amplification, *ALK* fusion, and WGD, each showing high activation, while other features have low activation scores across the WSI. These examples demonstrate the model’s specificity in capturing molecularly informative regions within histologically diverse tissue.

To further illustrate fine-grained spatial patterns, **Fig. 5** displays high-resolution Grad-CAM maps for four selected features: *TP53* mutation, *EGFR* p.L858R, WGD, and *KRAS* wild-type status. For *TP53, EGFR* p.L858R, and WGD (**Figs. 5a, 5b, and 5c**), high activation is consistently observed in tumor cells, while low activation score is observed in adjacent non-tumor cells. In contrast, **Fig. 5d** shows a tumor with mucinous histology and *KRAS* wild-type status, where low activation is observed throughout the WSI, consistent with the absence of clear morphologic correlates for this molecular profile.

**Fig. 5.**
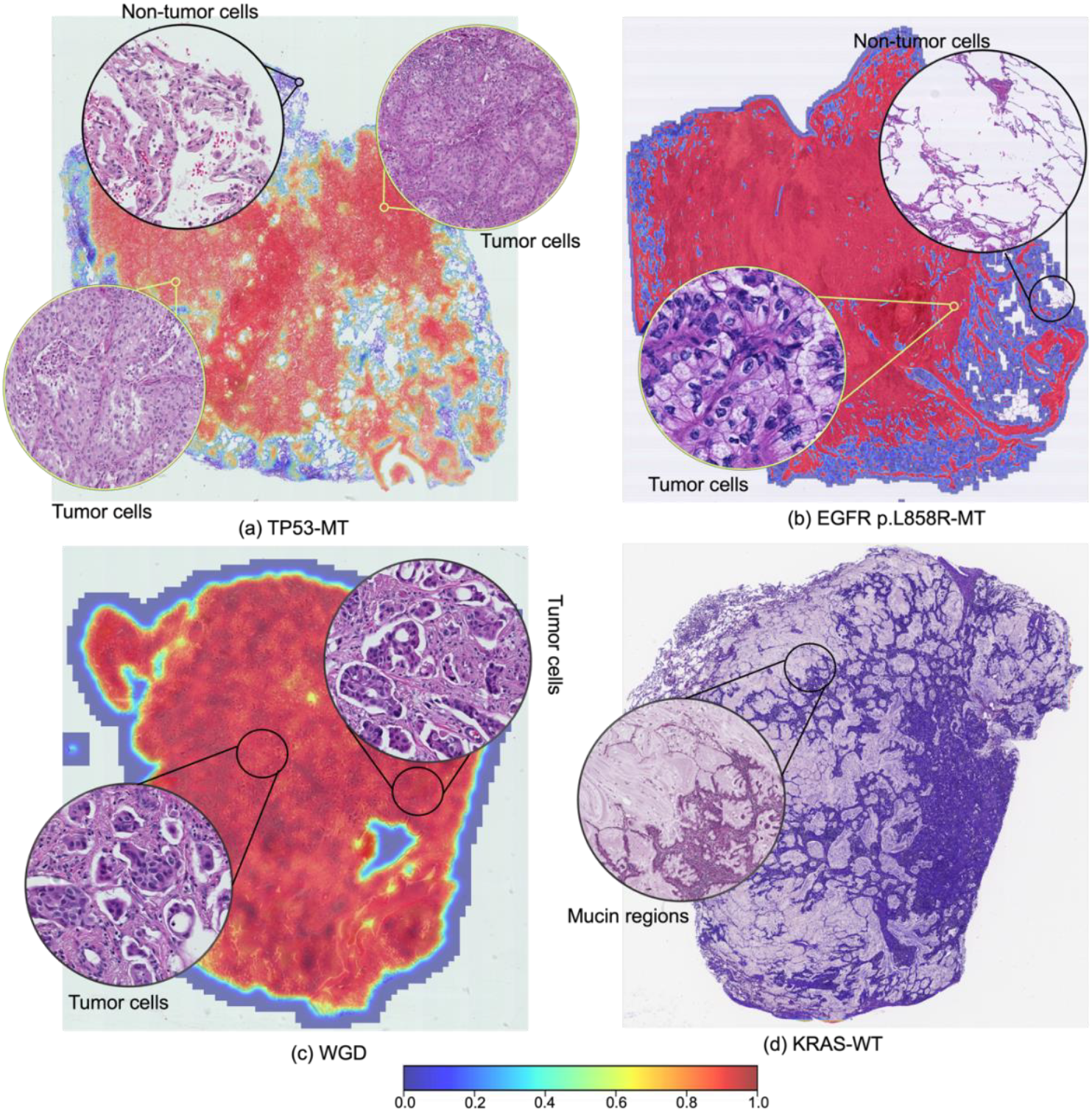
Zoomed-in activation maps highlighting spatial correlates of predicted molecular features. Representative Grad-CAM heatmaps for tumors predicted to harbor (a) *TP53* mutation, (b) *EGFR* p.L858R mutation, (c) whole-genome doubling, and (d) *KRAS* wild-type status, overlaid on H&E-stained whole slide images. Insets show magnified regions of interest with corresponding histologic features. The color bar indicates the intensity score. Higher intensity indicates a greater probability of being a region enriched with mutations (*TP53, EGFR, KRAS*) or additional molecular features (whole-genome doubling). MT: Mutant, WGD: Whole-genome doubling, WT: Wild-type

Together, these visualizations indicate that the model is capable of accurately detect spatially distinct histopathological signatures associated with specific molecular alterations, while minimizing false-positive signals in benign or non-informative regions. All example WSIs and corresponding heatmaps are publicly accessible via a public interactive web viewer developed in-house (WSI Image Viewer; **Supplementary Fig. 1**; see Code Availability).

## Discussion

Current standard-of-care molecular testing methods typically performed through NGS, while highly informative, remain expensive, time-consuming, and require access to specialized laboratory and bioinformatics infrastructure. These constraints can delay clinical decision-making and limit the widespread implementation of precision oncology, particularly in low-resource settings. In this study, we demonstrate that deep learning models, when applied to routine H&E-stained WSIs, can accurately discriminate a broad spectrum of genomic alterations in NS-LUAD. Using a weakly supervised, annotation-free, customized multilabel CNN, we achieved high predictive performance (AUROC ≥ 0.80) for 11 molecular alterations. These include not only typically tested *EGFR, KRAS*, and *ALK* overall alterations, but also genotype-specific mutations (e.g., *EGFR* p.L858R, p.E746_A750del), structural changes (e.g., *MDM2* amplification, *CDKN2A* deletion), and genome-wide features, such as WGD and kataegis. Although some of these genomic alterations are not currently targetable, they are strongly associated with tumor progression, therapeutic resistance, or poor prognosis [33-37] and could potentially inform clinical risk stratification and management. Moreover, our approach can identify molecular alterations in heterogenous tumors carrying multiple types of genomic alterations at the same time.

To our knowledge, this is the first deep learning model specifically developed for lung adenocarcinomas in subjects who have never smoked, an increasingly recognized lung cancer subtype with unique molecular and histologic characteristics. Our models outperformed the prior commonly used Inception-V3 [15] architecture across multiple key targets (i.e., *EGFR, KRAS, RBM10, EGFR* p.L858R, *EGFR* p.E476_A750del, *MDM2* amplification, and *ALK* fusion), and outperformed or showed similar detection capability to the standard ResNet50 model previously used to detect a limited number of genomic alterations [13, 26], which was the basis for our approach. These results highlight the importance and advantages of tailored model design for histology-based molecular prediction in NS-LUAD.

Importantly, we trained our model on the large, diverse, multi-institutional Sherlock-*Lung* cohort, capturing broad variability in tissue preparation, scanner types, and patient demographics. This design addresses longstanding concerns about the limited generalizability of models trained on homogeneous datasets, where performance differences may arise from batch effects, population differences, or both. By incorporating real-world heterogeneity, our study design improves model robustness and supports broader clinical translation. Moreover, our annotation-free framework—which includes both tumor and non-tumor regions—facilitates scalable integration into routine pathology workflows, where tumor purity is often variable and manual annotation is labor-intensive and subjective.

By uncovering fine-grained histologic–genomic relationships, our study also underscores the potential of morphology-based deep learning to bridge pathology and genomics, suggesting that spatially distinct histologic features reflect the aggregate phenotypic impact of underlying genetic alterations. To increase interpretability for the morpho-genomic relationship and address the “black box” nature of deep learning, we developed an interactive web-based platform that allows for a visual comparison of the model’s attention heatmaps against the original WSIs (**Supplementary Fig. 1**) [32], enabling pathologists to review regions contributing to model predictions. This freely available web tool (see **Code Availability** for the URL) was built to facilitate the identification and validation of spatial patterns that the model associates with specific genomic alterations. Such interpretability is essential for clinical adoption, enabling collaborative human–AI decision-making, and bridging the gap between automated inference and expert interpretation and supporting trust among clinicians and patients [38].

Despite these strengths, our study has limitations. Rare alterations in NS-LUAD, such as *KRAS* p.G12C, *KRAS* p.G12D, *KRAS* p.G12V, *RBM10* mutations, and *ALK* fusions were underrepresented (prevalence <10%) in our dataset, potentially limiting predictive performance. Although we employed data augmentation strategies to address the low prevalences, future work should incorporate larger datasets as well as explore transfer learning and transformer-based approaches to improve accuracy for low-prevalence mutations. This further optimization is needed before clinical integration, particularly for triage applications, where high sensitivity is critical to minimize false negatives and avoid missing therapeutic opportunities [39].

Additionally, some alterations, such as the APOBEC signature remained challenging to predict using both our customized multilabel CNN and other established architectures, suggesting that not all mutations manifest morphologically in a consistent or detectable manner [40-42]. Integrating histologic models with orthogonal modalities such as spatial transcriptomics [43, 44] or multiplex immunofluorescence [45] could provide deeper molecular context for AI-derived predictions. These multilabel approaches hold promise for refining histology-based models and improving their biological interpretability by elucidating how genetic alterations shape tissue architecture and the tumor microenvironment [46].

In conclusion, we developed a generalizable, annotation-free deep learning model that accurately predicts a broad range of genomic alterations from routine histology in NS-LUAD. Our approach offers a cost-effective, scalable solution to augment molecular diagnostics. This is particularly relevant in resource-constrained settings, where it may accelerate tissue triaging, enhance clinical decision-making, reduce diagnostic delays, and expand access to precision oncology.

## Author Contribution

**Conceptualization:** M.S., T.Z., S.-R.Y., J.S.A., and M.T.L.; **Manuscript original draft:** M.S., T.-V.-T.T., and M.T.L.; **Data access and curation:** M.S., T.Z., W.Z., P.H.H., K.M., S.M.L., N. R., and Q. L.; **Formal analysis and methodology development:** M.S.; **Software development supervision:** J.S.A.; **Software implementation:** M.S., P.M.S.B., and J.S.A.; **Pathology evaluation:** R.H., M.K.B., L.M.S., P.J., C.L., W.D.T., and S.-R.Y.; **Statistical support:** J.S.; **Resources:** S.J.C. and M.T.L.; **Study supervision:** M.T.L. All authors reviewed the manuscript.

## Ethics Statement and Patient Consent

Because the National Cancer Institute (NCI) only received de-identified samples and data from collaborating centers, had no direct contact or interaction with the study participants and did not use or generate identifiable private information, individual patient consent was not required by the NCI. As a result, the Sherlock-Lung study has been determined to constitute “NotHuman Subject Research (NHSR)” under the Federal Common Rule (45 CFR 46; https://www.ecfr.gov/), with all data collected in accordance with locally approved Institutional Review Board protocols.

## Funding

This research was supported by the Intramural Research Program of the National Institutes of Health (NIH) (project ZIACP101231 to MTL) and the Anne Wojcicki Foundation (Grant Number: LC009). The contributions of the NIH author(s) were made as part of their official duties as NIH federal employees, are in compliance with agency policy requirements, and are considered Works of the United States Government. However, the findings and conclusions presented in this paper are those of the author(s) and do not necessarily reflect the views of the NIH or the U.S. Department of Health and Human Services.

## Acknowledgements

All analyses were performed using the computational resources of the NIH Biowulf high-performance computing cluster (https://hpc.nih.gov/).

## Data availability

Whole-genome sequencing tumor CRAM files from the Sherlock-*Lung* study used in this analysis have been deposited in dbGaP under accession number phs001697.v2.p1 (available at: https://www.ncbi.nlm.nih.gov/gap/advanced_search/?TERM=phs001697.v2.p1).

## Code Availability

The code for this study is accessible on GitHub: https://github.com/monjoybme/Mutation_AI. An in-house web-based tool (Sherlock WSI Viewer) was developed to facilitate the real-time use of the CNN activation heatmaps by the pathologist as a “digital stain” on the WSI. The tool is freely available at: https://episphere.github.io/sherlockWSI/#gcsBaseFolder=301f394c-ea68-4a26-829b-88208fffb5c6. The Sherlock WSI Viewer displays the output of our customized multilabel CNN in the form of heatmaps overlaid on the original H&E-stained whole slide images, representing 16 molecular characteristics that can be selected individually from a dropdown menu (left corner). The right panel presents the corresponding original H&E-stained whole slide image. In the heatmap view, red regions indicate high activation scores, while blue regions indicate low activation scores. Users can switch between WSIs using the “Image Name” dropdown menu located at the top center of the interface. For additional details on activation scores and molecular characteristics, refer to **Fig. 4**.

## Notes

### Competing Interest Statement

SRY has received consulting fees from AstraZeneca, Sanofi, Amgen, AbbVie, and Sanofi; received speaking fees from AstraZeneca, Medscape, PRIME Education, and Medical Learning Institute. All other authors declare that they have no competing interests.

### Summary of Updates

This version of the manuscript has been revised to include the following updates: the title has been updated, the affiliation of Dr. Landi has been revised, and the Anne Wojcicki Foundation has been added as a new funder. In addition, the title page has been removed from the supplementary materials.

## References

1. Khan, S., et al., Lung cancer in never smokers (LCINS): development of a UK national research strategy. BJC Reportss, 2023. 2(1): p. 1.

2. LoPiccolo, J., et al., Lung cancer in patients who have never smoked—An emerging disease. Nature Reviews Clinical Oncology, 2024. 21(2): p. 121–146.

3. Chapman, A.M., et al., Lung cancer mutation profile of EGFR, ALK, and KRAS: Meta-analysis and comparison of never and ever smokers. Lung Cancer, 2016. 102: p. 122–134.

4. Zhang, T., et al., Genomic and evolutionary classification of lung cancer in never smokers. Nat Genet, 2021. 53(9): p. 1348–1359.

5. Jaaks, P., et al., Effective drug combinations in breast, colon and pancreatic cancer cells. Nature, 2022. 603(7899): p. 166–173.

6. Meric-Bernstam, F., et al., National Cancer Institute Combination Therapy Platform Trial with Molecular Analysis for Therapy Choice (ComboMATCH). Clin Cancer Res, 2023. 29(8): p. 1412–1422.

7. Echle, A., et al., Deep learning in cancer pathology: a new generation of clinical biomarkers. British Journal of Cancer, 2021. 124(4): p. 686–696.

8. He, K., et al. Deep residual learning for image recognition. in Proceedings of the IEEE conference on computer vision and pattern recognition. 2016.

9. Szegedy, C., et al. Rethinking the inception architecture for computer vision. in Proceedings of the IEEE conference on computer vision and pattern recognition. 2016.

10. Fu, Y., et al., Pan-cancer computational histopathology reveals mutations, tumor composition and prognosis. Nat Cancer, 2020. 1(8): p. 800–810.

11. Kather, J.N., et al., Pan-cancer image-based detection of clinically actionable genetic alterations. Nat Cancer, 2020. 1(8): p. 789–799.

12. Tomita, N., et al., Predicting oncogene mutations of lung cancer using deep learning and histopathologic features on whole-slide images. Translational Oncology, 2022. 24: p. 101494.

13. Pao, J.J., et al., Predicting EGFR mutational status from pathology images using a real-world dataset. Sci Rep, 2023. 13(1): p. 4404.

14. Jain, M.S. and T.F. Massoud, Predicting tumour mutational burden from histopathological images using multiscale deep learning. Nature Machine Intelligence, 2020. 2(6): p. 356–362.

15. Coudray, N., et al., Classification and mutation prediction from non-small cell lung cancer histopathology images using deep learning. Nat Med, 2018. 24(10): p. 1559–1567.

16. Zhao, Y., et al., Deep learning using histological images for gene mutation prediction in lung cancer: a multicentre retrospective study. Lancet Oncol, 2025. 26(1): p. 136–146.

17. Campanella, G., et al., Real-world deployment of a fine-tuned pathology foundation model for lung cancer biomarker detection. Nat Med, 2025.

18. Terada, Y., et al., Artificial Intelligence-Powered Prediction of ALK Gene Rearrangement in Patients With Non-Small-Cell Lung Cancer. JCO Clin Cancer Inform, 2022. 6: p. e2200070.

19. Landi, M.T., et al., Tracing Lung Cancer Risk Factors Through Mutational Signatures in Never-Smokers: The Sherlock-Lung Study. American Journal of Epidemiology, 2021. 190(6): p. 962–976.

20. Zhang, T., et al., Deciphering lung adenocarcinoma evolution and the role of LINE-1 retrotransposition. bioRxiv, 2025: p. 2025.03. 14.643063.

21. Moreira, A.L., et al., A Grading System for Invasive Pulmonary Adenocarcinoma: A Proposal From the International Association for the Study of Lung Cancer Pathology Committee. J Thorac Oncol, 2020. 15(10): p. 1599–1610.

22. Diaz-Gay, M., et al., The mutagenic forces shaping the genomes of lung cancer in never smokers. Nature, 2025. 644(8075): p. 133–144.

23. Jongjoo, K., S.K. Davis, and J.F. Taylor, Application of non-parametric bootstrap methods to estimate confidence intervals for QTL location in a beef cattle QTL experimental population. Genet Res, 2002. 79(3): p. 259–63.

24. Liu, Y., M.R. Smith, and R.M. Rangayyan, The application of Efron’s bootstrap methods in validating feature classification using artificial neural networks for the analysis of mammographic masses. Conf Proc IEEE Eng Med Biol Soc, 2004. 2004: p. 1553–6.

25. Youden, W.J., Index for rating diagnostic tests. Cancer, 1950. 3(1): p. 32–5.

26. Zhao, D., et al., High accuracy epidermal growth factor receptor mutation prediction via histopathological deep learning. BMC Pulm Med, 2023. 23(1): p. 244.

27. DeLong, E.R., D.M. DeLong, and D.L. Clarke-Pearson, Comparing the areas under two or more correlated receiver operating characteristic curves: a nonparametric approach. Biometrics, 1988. 44(3): p. 837–45.

28. Benjamini, Y. and Y. Hochberg, Controlling the False Discovery Rate - a Practical and Powerful Approach to Multiple Testing. Journal of the Royal Statistical Society Series B-Statistical Methodology, 1995. 57(1): p. 289–300.

29. Selvaraju, R.R., et al. Grad-cam: Visual explanations from deep networks via gradient-based localization. in Proceedings of the IEEE international conference on computer vision. 2017.

30. Selvaraju, R.R., et al., Grad-CAM: Visual Explanations from Deep Networks via Gradient-Based Localization. International Journal of Computer Vision, 2020. 128(2): p. 336–359.

31. contributors, O. OpenSeadragon: An open-source, web-based viewer for high-resolution images. [cited 2025 June 30]; Available from: https://openseadragon.github.io.

32. Bhawsar, P., et al., ImageBox3: No-Server Tile Serving to Traverse Whole Slide Images on the Web. arXiv preprint 2207.01734, 2022.

33. Sinha, A., et al., Early-Stage Lung Adenocarcinoma MDM2 Genomic Amplification Predicts Clinical Outcome and Response to Targeted Therapy. Cancers (Basel), 2022. 14(3).

34. Gutiontov, S.I., et al., CDKN2A loss-of-function predicts immunotherapy resistance in non-small cell lung cancer. Sci Rep, 2021. 11(1): p. 20059.

35. Reyes, A., et al., RBM10 Mutation as a Potential Negative Prognostic/Predictive Biomarker to Therapy in Non-Small-Cell Lung Cancer. Clin Lung Cancer, 2024. 25(8): p. e411–e419.

36. Zhao, J., et al., Functional analysis reveals that RBM10 mutations contribute to lung adenocarcinoma pathogenesis by deregulating splicing. Sci Rep, 2017. 7: p. 40488.

37. Lopez, S., et al., Interplay between whole-genome doubling and the accumulation of deleterious alterations in cancer evolution. Nat Genet, 2020. 52(3): p. 283–293.

38. Ghassemi, M., L. Oakden-Rayner, and A.L. Beam, The false hope of current approaches to explainable artificial intelligence in health care. Lancet Digit Health, 2021. 3(11): p. e745–e750.

39. Power, M., G. Fell, and M. Wright, Principles for high-quality, high-value testing. Evid Based Med, 2013. 18(1): p. 5–10.

40. Tamura, R., et al., Spatial genomic diversity associated with APOBEC mutagenesis in squamous cell carcinoma arising from ovarian teratoma. Cancer Sci, 2023. 114(5): p. 2145–2157.

41. Roper, N., et al., APOBEC Mutagenesis and Copy-Number Alterations Are Drivers of Proteogenomic Tumor Evolution and Heterogeneity in Metastatic Thoracic Tumors. Cell Rep, 2019. 26(10): p. 2651–2666 e6.

42. Roberts, S.A., et al., An APOBEC cytidine deaminase mutagenesis pattern is widespread in human cancers. Nat Genet, 2013. 45(9): p. 970–6.

43. Ge, Y., et al., Deep Learning-Enabled Integration of Histology and Transcriptomics for Tissue Spatial Profile Analysis. Research (Wash D C), 2025. 8: p. 0568.

44. Zeng, Y., et al., Spatial transcriptomics prediction from histology jointly through Transformer and graph neural networks. Brief Bioinform, 2022. 23(5).

45. Wu, E., et al., ROSIE: AI generation of multiplex immunofluorescence staining from histopathology images. bioRxiv, 2024.

46. Jiang, X., et al., iIMPACT: integrating image and molecular profiles for spatial transcriptomics analysis. Genome Biol, 2024. 25(1): p. 147.

