## Supplementary Material for "Genomic Characterization of Lung Cancer in Never-Smokers Using Deep Learning"

***Abbreviations***:

- **EGFR**: Epidermal Growth Factor Receptor
- **KRAS**: Kirsten Rat Sarcoma Viral Oncogene Homolog
- **TP53**: Tumor Protein p53
- **LRP1B**: Low-Density Lipoprotein Receptor-Related Protein 1B
- **STK11**: Serine/Threonine Kinase 11
- **TMB**: Tumor Mutational Burden
- **CDKN2A**: Cyclin-Dependent Kinase Inhibitor 2A
- **MDM2**: Mouse Double Minute 2 Homolog
- **ALK**: Anaplastic Lymphoma Kinase
- **RBM10**: RNA Binding Motif Protein 10
- **APOBEC**: Apolipoprotein B mRNA Editing Enzyme, Catalytic Polypeptide-like
- **WGD**: Whole-Genome Doubling

**EGFR Mutations:**

1. **p.L858R**: Substitution of leucine (L) with arginine (R) at position 858 in the EGFR protein.
2. **p.E746_A750del**: Deletion of amino acids from glutamic acid (E) at position 746 to alanine (A) at position 750 in the EGFR protein.

**KRAS Mutations:**

1. **p.G12C**: Substitution of glycine (G) with cysteine (C) at position 12 in the KRAS protein.
2. **p.G12V**: Substitution of glycine (G) with valine (V) at position 12 in the KRAS protein.
3. **p.G12D**: Substitution of glycine (G) with aspartic acid (D) at position 12 in the KRAS protein.

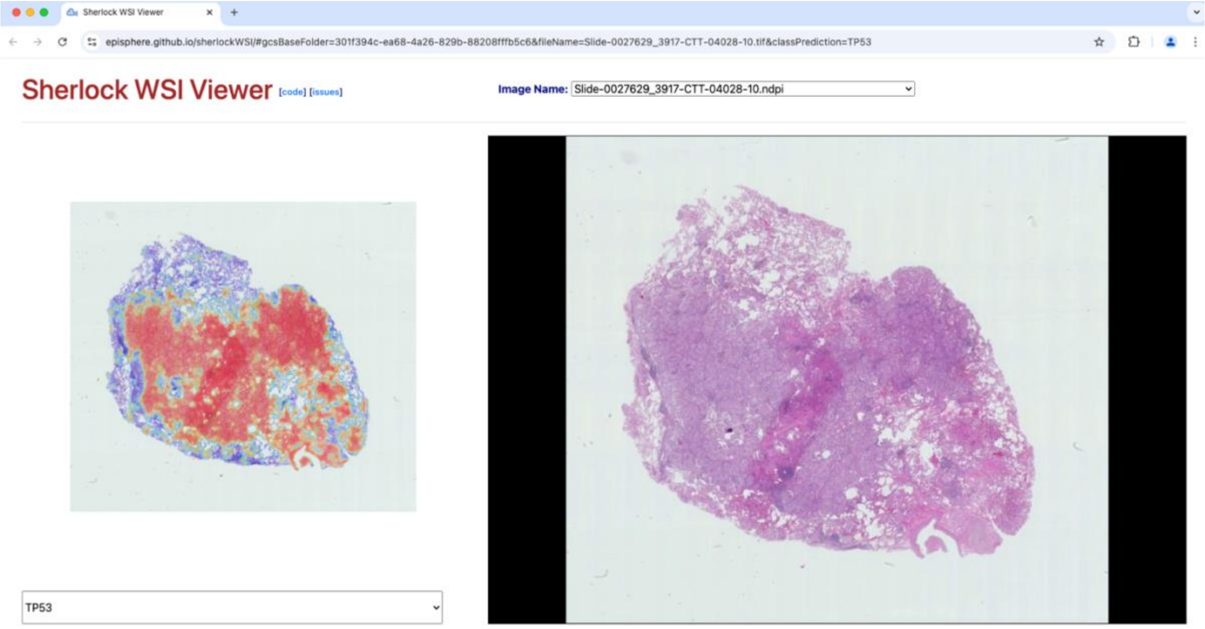


**Supplementary Fig. 1**: Screenshot of the interactive web-based tool displaying convolutional neural network activation heatmaps for a representative case with TP53 mutation in a lung adenocarcinoma from an individual who has never smoked (The tool is freely available at: https://episphere.github.io/sherlockWSI/#gcsBaseFolder=301f394c-ea68-4a26-829b-88208fffb5c6).
